# Salience-dependent disruption of sustained auditory attention can be inferred from evoked pupil responses and neural tracking of task-irrelevant sounds

**DOI:** 10.1101/2023.10.30.564684

**Authors:** Lorenz Fiedler, Ingrid Johnsrude, Dorothea Wendt

**Author notes:** Corresponding Author: Lorenz Fiedler.

## Abstract

Stimulus-driven attention allows us to react to relevant stimuli (and imminent danger!) outside our current focus of attention. But irrelevant stimuli can also disrupt attention; for example, during listening to speech. The degree to which sound captures attention is called salience, which can be estimated by existing, behaviorally validated, computational models (Huang & Elhilali, 2017). Here we examined whether neurophysiological responses to task-irrelevant sounds indicate the degree of distraction during a sustained-listening task and how much this depends on individual hearing thresholds. N = 47 Danish-speaking adults (28/19 female/male; mean age: 60.1, SD: 15.9 years) with heterogenous hearing thresholds (PTA; mean: 25.5, SD: 18.0 dbHL) listened to continuous speech while one-second-long, task-irrelevant natural sounds (distractors) of varying computed salience were presented at unpredictable times and locations. Eye tracking and electroencephalography were used to estimate pupil response and neural tracking, respectively. The task-irrelevant sounds evoked a consistent pupil response (PR), distractor-tracking (DT) and a drop of target-tracking (ΔTT), and statistical modelling of these three measures within subjects showed that all three are enhanced for sounds with higher computed salience. Participants with larger PR showed a stronger drop in target tracking (ΔTT) and performed worse in target speech comprehension. We conclude that distraction can be inferred from neurophysiological responses to task-irrelevant stimuli. These results are a first step towards neurophysiological assessment of attention dynamics during continuous listening, with potential applications in hearing-care diagnostics.

**Significance statement:** Successful speech-in-noise comprehension in daily life does not only depend on the acuity of the auditory input, but also cognitive factors like attentional control. Being able to measure distraction-dependent neurophysiological responses to peripheral, task-irrelevant stimuli would enable monitoring the extent to which the attentional focus is instantaneously captured away from a target under sustained attention. Here we show that especially pupil response and neural tracking of distractor sounds reflect the degree to which people with both normal and elevated hearing thresholds are distracted. Such a measure could be used to non-invasively track the focus of attention and thus could find application in hearing care diagnostics, where cognitive factors like attentional control are being increasingly recognized as important.

## Introduction

Distractibility is a vital feature of our attentional system. For example, we still need to hear a car approaching while we focus our attention on the traffic light across the street. While voluntary attention (a.k.a. top-down attention, intentional attention) narrows down incoming information to a minimum, peripheral stimuli can still capture our attention in a stimulus-driven fashion (also called automatic or bottom-up attention). Unfortunately, some sounds may capture our attention for no relevant reason. Hearing comes with the vital blessing of helping us to detect potential danger from any direction, but with the curse of leading to distraction by irrelevant sounds.

The degree to which a sound captures stimulus-driven attention is called salience (Kaya & Elhilali, 2014). Voluntary selective auditory attention modulates neural response mainly in cortex (Mesgarani & Chang, 2012), although recent studies appear to demonstrate attentional modulation more peripherally in the brainstem (Forte et al., 2017) and cochlea (Gehmacher et al., 2022). How, and where, stimulus-driven attention is implemented along the auditory pathway and which stimulus features are driving it is as yet unclear (Huang & Elhilali, 2020; Serences & Kastner, 2014). For example, a sound can capture attention due to its sheer loudness independent of its meaning, whereas our own name may capture our attention even if it is spoken at a moderate level (Cherry, 1953). Here we first focus on objective, acoustic features that give rise to salience and second, the beyond-acoustic aspects of salience that may be captured in the statistical dependencies in the neurophysiological measures and their ability to predict behavior.

There have been several attempts to create models for auditory salience. Inspired by vision research, earlier attempts (Duangudom & Anderson, 2007; Kalinli & Narayanan, 2007; Kaya & Elhilali, 2012; Kayser et al., 2005) treated the spectrogram like a picture and thus have been criticized for ignoring the unfolding of a sound in time (Kaya & Elhilali, 2017). Later, the temporal unfolding of established psychoacoustic features such as loudness, harmonicity and pitch were incorporated into the models (Huang & Elhilali, 2017; Kaya & Elhilali, 2014). Such models are meant to explain an automatic, stimulus-driven behavior, but have been validated behaviorally using explicit tasks, such as rating and detection. The question is whether these models hold in continuous listening tasks with participants naïve about the purpose of the study. Only recently, these models were found to explain some variability in neural phase-locking during continuous listening paradigms (Huang & Elhilali, 2020; Straetmans et al., 2021). While these studies investigated event-related potentials to discrete events, we here aim to generalize their findings to cortical tracking of continuous stimuli and pupillometry.

Qualitatively, listeners with hearing impairment complain about a lack of situational awareness on the one hand, but also about distraction by irrelevant sounds on the other (Nielsen & Henriksen, 2022). That means they are lacking selectivity in the processing of salient stimuli, which may have two causes: 1) hearing loss leads to a degraded input to the auditory pathway which corrupts features essential for stream segregation and selection (David et al., 2018; Oxenham, 2008); 2) Increased cognitive load and fatigue may limit the resources available for attentional control (Peelle, 2018). To better understand why selectivity in perception of salient stimuli is reduced in people with HI, we need methods that can assess the degree of distraction in scenarios of high ecological validity such as continuous speech.

We asked if established neurophysiological measures for attention (cortical tracking) and cognitive, listening-related effort (pupil size, see methods) correlate with the output of a recent model for auditory salience (Huang & Elhilali, 2017, 2020). We examined the neurophysiological response to task-irrelevant sounds in participants with heterogenous hearing profiles. We hypothesize that: (1) there is a consistent neurophysiological response (i.e., neural tracking and pupil response) to task-irrelevant sounds; (2) these responses correlate positively with computed salience; (3) participants with increased responses perform worse in the sustained auditory attention task.

## Methods

### Participants

Our goal was to recruit a group of participants with heterogenous age and hearing loss profiles. A total of N = 47 participants (28 female) were enrolled in this study: 24 of them were hearing-aid users (with sensorineural hearing loss). The age of the participants ranged from 23 to 82 years (mean: 60.1, SD: 15.9). The pure-tone average thresholds (PTA; 500-4000 Hz), calculated from the better ear at each frequency, ranged from −0.7 to 52.2 dB HL (mean: 25.5, SD: 18.0). Figure 2A depicts hearing thresholds of the better ear and Figure 2B depicts the correlation between PTA and age. In the statistical modelling, we controlled for age effects before PTA was used as a predictor (see statistical modelling). The study was approved by the Copenhagen regional ethics committee: (De Videnskabsetiske Komitéer, Region Hovedstaden).

### Apparatus

The experiment took place in an acoustically and electro-magnetically shielded booth. Participants were seated in a comfortable office chair. The office chair was placed such that participants were seated at a reference point in the middle of the room and their ears were level with the loudspeakers. A computer keyboard was placed on a desk in front of the participants. Left and right Ctrl-buttons were labelled with “Ja” & “Nej” (Yes & No). The assignment of the labels to left and right was balanced across participants. The space bar was labelled with “Start”. Behind the keyboard, a Tobii Pro Spectrum eye tracker (Tobii AB, Stockholm, Sweden) was placed. The light conditions were kept at a moderate level and measured at 75 lux at the reference point.

Three loudspeakers (Genelec 8030A, Genelec Oy, Iisalmi, Finland) were placed at a radius of one meter from the reference point. One loudspeaker was positioned at 0 degrees azimuth (front, behind the desk, above the screen), and two were positioned to the left and right at ±90 degrees azimuth. The loudspeakers were connected to an RME Multiface II audio interface (AudioAG, Haimhausen, Germany). The audio interface was connected via USB to a personal computer running Microsoft Windows 10 (Microsoft Corp., Redmond, USA), where MATLAB 2020b (Mathworks Inc., Natick, USA) and Tobii Eye Tracker Manager were installed. The audio was controlled from MATLAB using Soundmexpro toolbox (Hörtech gGmbH, Oldenburg, Germany). Underneath the frontal loudspeaker, a screen was placed to display task instructions and comprehension questions. The screen was controlled via MATLAB using Psychtoolbox version 3.0.17 (Brainard, 1997).

A BioSemi ActiveTwo EEG amplifier (BioSemi B.V., Amsterdam, The Netherlands) with 64 channels was placed behind the participant. A BioSemi USB Trigger Interface cable connected the computer with the BioSemi receiver. This connection was used to send timing markers, while precise timing was ensured by sending triggers to the EEG amplifier via StimTrak (Brain Products GmbH, Gilching, Germany) from the audio interface. For EEG data collection, the BioSemi receiver was connected via USB to another, identical computer, which had BioSemi ActiView installed.

### Stimulus material

#### Target stimuli

A collection of 320 continuous speech recordings, each with a length of 30 seconds, were used as target stimuli. The recordings were local radio newsclips (used in earlier studies Alickovic et al., 2020, 2021; Fiedler et al., 2021) spoken by four different Danish radio announcers (2 female, 2 male). The root-mean-square (RMS) of the recordings was equalized, such that each of the 30-second-long recordings had the same RMS.

#### Distractor sounds

The goal was to collect sounds that varied in spectro-temporal profiles. A collection of 690, one-second-long natural sounds was compiled from freely available sources such as freesound.org, mixkit.co, as well as existing databases from former studies (Ananthabhotla et al., 2019; Gygi & Shafiro, 2019; Santoro et al., 2014; Zhao et al., 2019). The collection consisted of sounds from animals, tools, machines, music, human sounds like coughing and laughing, but no speech. The RMS of the distractor sounds was equalized, such that each of the one-second-long sounds had the same RMS. Subsequently, the overall level of the sounds was lowered by 10 dB, such that the RMS of the distractor sounds was 10 dB below the RMS of the target stimuli.

The distractor sounds were embedded in stationary noise to keep the masking of the target speech constant in a long-term sense (Fig. 1B). This way, we made sure that distraction is not mainly driven by increased energetic but rather informational masking. To this end, white noise was created and filtered such that its long-term average spectrum (LTAS) fitted the LTAS of all distractor sounds following procedures of May et al. (2018). In brief, the LTAS of all concatenated distractor stimuli was calculated in bandwidths of 1/3 octaves. A finite-impulse response filter was designed (using the function fir2 in MATLAB) and applied to the white noise such that its LTAS was adjusted to match the LTAS of the distractor sounds.

**Figure 1:**
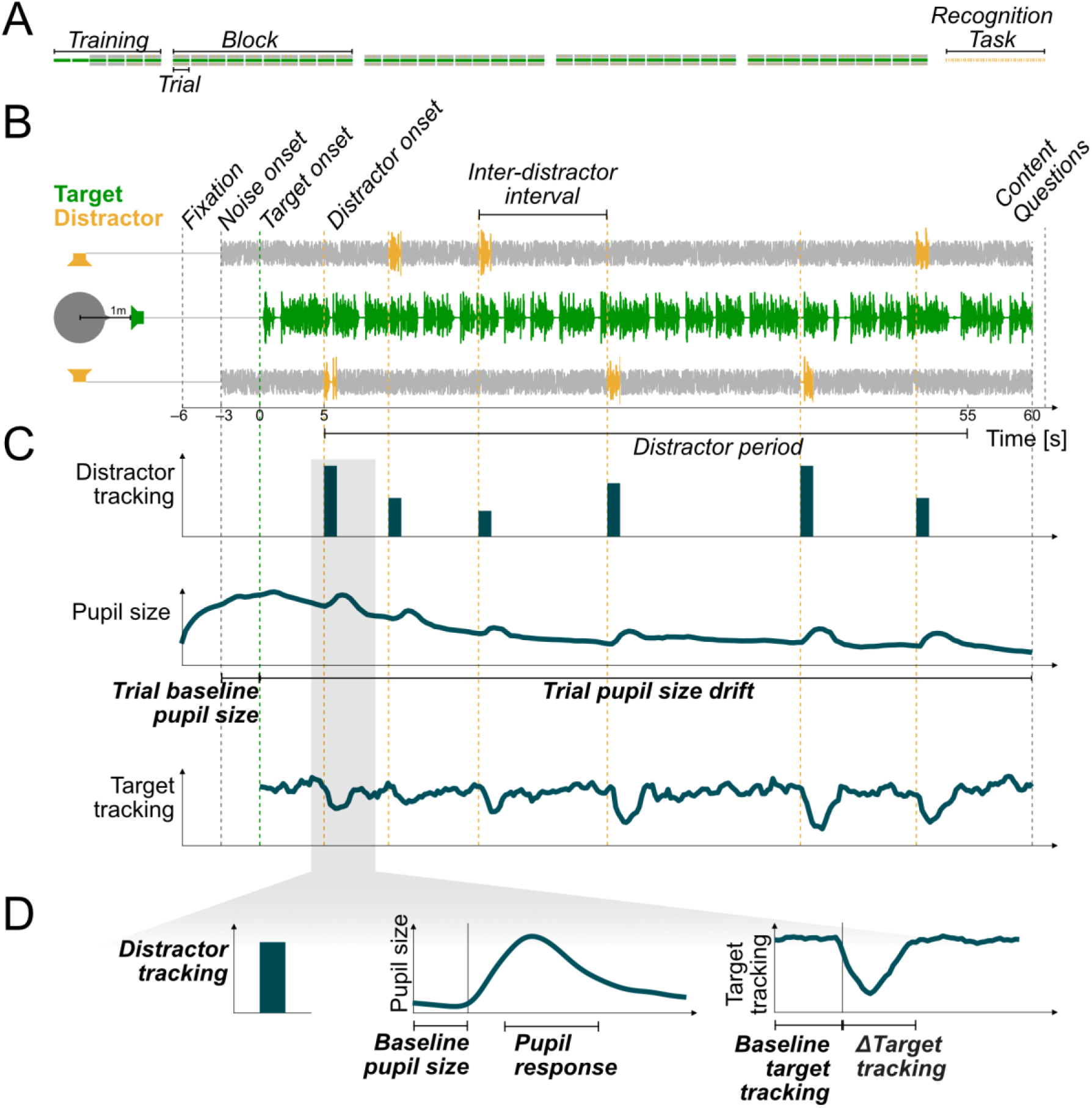
Study design. Neurophysiological covariates and measures of interest are depicted in bold italic. **A)** The main experiment consisted of 46 trials (the first 6 trials were training trials) followed by a surprise recognition task. **B)** Three loudspeakers were set up at the respective angles of 0 and ±90 degrees. Each trial started with three seconds of silence, followed by the onset of stationary noise from the loudspeakers at ±90 degrees. Three seconds later, target speech was presented from the loudspeaker at 0 degrees. Between 5 seconds post target onset (earliest) and 55 seconds post target onset (latest), 1-second-long sounds were presented randomly at + or – 90 degrees and with a jittered onset interval between 5 and 10 seconds. **C)** Trial-related measures (time rel. to speech onset): Distractor tracking was directly derived from the one-second presentation window of the distractor embedded in noise. Pupil size: Covariates trial baseline pupil size (mean −3 to 0 sec) and trial pupil size drift (mean 0 to 60 sec). Continuous target tracking was derived by sliding a window of one-second width in steps of 0.1 seconds. **D)** Distractor-related measures (time rel. to distractor onset): Distractor tracking (see C), covariate baseline pupil size (mean −1 to 0 sec), pupil response (mean change from baseline 1 to 3 sec), covariate baseline target tracking (derived from a window between −2 and 0 sec), and Δtarget tracking (change from baseline derived from a window between 0 and 2 sec).

#### Randomization

We randomized the selection of target stimuli such that, for each participant, 92 of the 320 30-sec target stimuli were selected and one was appended to the first to make a one-minute stimulus (Figure 1B). Each participant heard 46 such one-minute-long stimuli, one on each trial. The concatenated stimuli always consisted of two different target talkers and after 30 seconds there was a switch of target talker. While we deliberately presented different target talkers to make the results more generalizable, the switch in target talkers after 30 seconds was not of particular interest to us but was a by-product of our aim to create one-minute long target stimuli.

We randomized the selection of distractor stimuli by randomly selecting 280 out of the 690 different sounds for each participant. An inter-distractor interval for distractor onsets was randomly assigned to each of the distractor stimuli by drawing from a discrete uniform distribution between 5 and 10 seconds in steps of 0.2 seconds. Furthermore, the distractor location “left” or “right” was randomly assigned with the exception that the same location was never repeated more than 5 times in a row.

The reasoning behind presenting different stimuli to different participants was to maximize the generalizability of the results and thus allow later classification approaches on the same data set by avoiding the risk of overfitting. For the analysis presented here we consider the variability in stimuli neither necessary nor limiting.

Based on the randomization of target and distractor stimuli, individual three-channel audio files were created for each trial (Figure 1B). Each sound file started with three seconds of silence, after which the noise started at the left and right loudspeakers (±90°). To ensure that the two noise sources were uncorrelated, different sections of the noise were presented on the two loudspeakers. After 3s of noise, the target stimulus started at the loudspeaker in front (0°).

The distractor sounds were presented at the left and right loudspeakers embedded in the noise (Figure 1B). Within the one-minute-long trials, distractor stimuli could occur between 5 and 55 seconds relative to target onset (the ‘distractor period’). This was done to ensure that there was enough time for the participant to establish an attentional focus on the target talker at the beginning of the trial and that there was enough time to collect the physiological responses to the last distractor sound of the trial.

To uniformly distribute the distractor sounds across the distractor period, the inter-distractor interval was carried over from one trial to the next. For example, if the last distractor in trial n-1 was presented at 51 seconds, and the next scheduled distractor had an associated inter-distractor interval of 8 seconds, 4 seconds of that were used at the end of trial n-1 (i.e., between 51 and 55 sec), and the remaining 4 seconds were appended at the beginning of trial n (i.e., next distractor started 5 sec plus 4 sec = 9 sec into trial n). At the beginning of each distractor sound, the noise was faded out, and was faded back in again at the end of the distractor using 20-msec half-sinusoidal ramps. Since the noise had the same LTAS as the distractors, the fade-out/fade-in ensured that, during the distractor sounds, the energetic masking was constant in the long-term average sense.

The experiment was blocked, and participants could take breaks between blocks. The first six trials were for training (Figure 1A). In the first two trials, only the target stimuli were presented. The stationary noise was introduced in trials 3 & 4 and the distractors were introduced in trials 5 & 6. Forty trials were presented in the main experiment. Together with the two last trials from the training, 42 trials were presented with distractor sounds; with largely different targets and distractors for each participant, as described above.

### Hearing loss compensation and adaptation

Hearing-aid users were not wearing their hearing aids during the experiment. To make sure that all participants were presented with stimuli of comparable loudness, we individualized the amplification in two steps (as described in detail below): first, we used linear gain to spectrally equalize (i.e., flatten) local dips in their pure-tone audiograms and second, we estimated a stimulus-specific hearing threshold. The resulting amplification was applied to the stimuli before presentation.

The linear gain was applied by designing an individual finite-impulse response (FIR) filter for each participant, with the goal to compensate local dips in the audiogram. We aimed to provide gain to frequencies that exceeded 20 dB difference to the best hearing threshold. The presentation in free field only allowed for one, better-ear gain profile to make sure that we would not present at uncomfortable, excessively loud, levels. First, we measured the individual pure-tone audiograms to identify, for each person, for each frequency (125, 250, 750, 1000, 1500, 2000, 3000, 4000, 6000, 8000 Hz) the lower (i.e., better) hearing threshold (HT) between the two ears (Figure 2A). Second, we identified the lowest (i.e., best) HT across these measurements, which can be seen as a reference peak in the audiogram. Third, we calculated the to-be-added gain for frequencies with an HT more than 20 dB higher (i.e., worse) than the reference. The gain was calculated from the difference between the specific HT and the best HT minus 20 dB. Hence, we filled up dips in the audiogram that exceeded 20 dB relative to the reference peak. To derive the gain for a 1/3-octave spaced FIR-filter, we took the gain calculated at the frequencies of the audiogram as support points and interpolated the gain values missing on a 1/3-octave-spaced frequency axis. Finally, an FIR-filter was designed using the function fir2 in MATLAB and applied to all stimuli.

**Figure 2:**
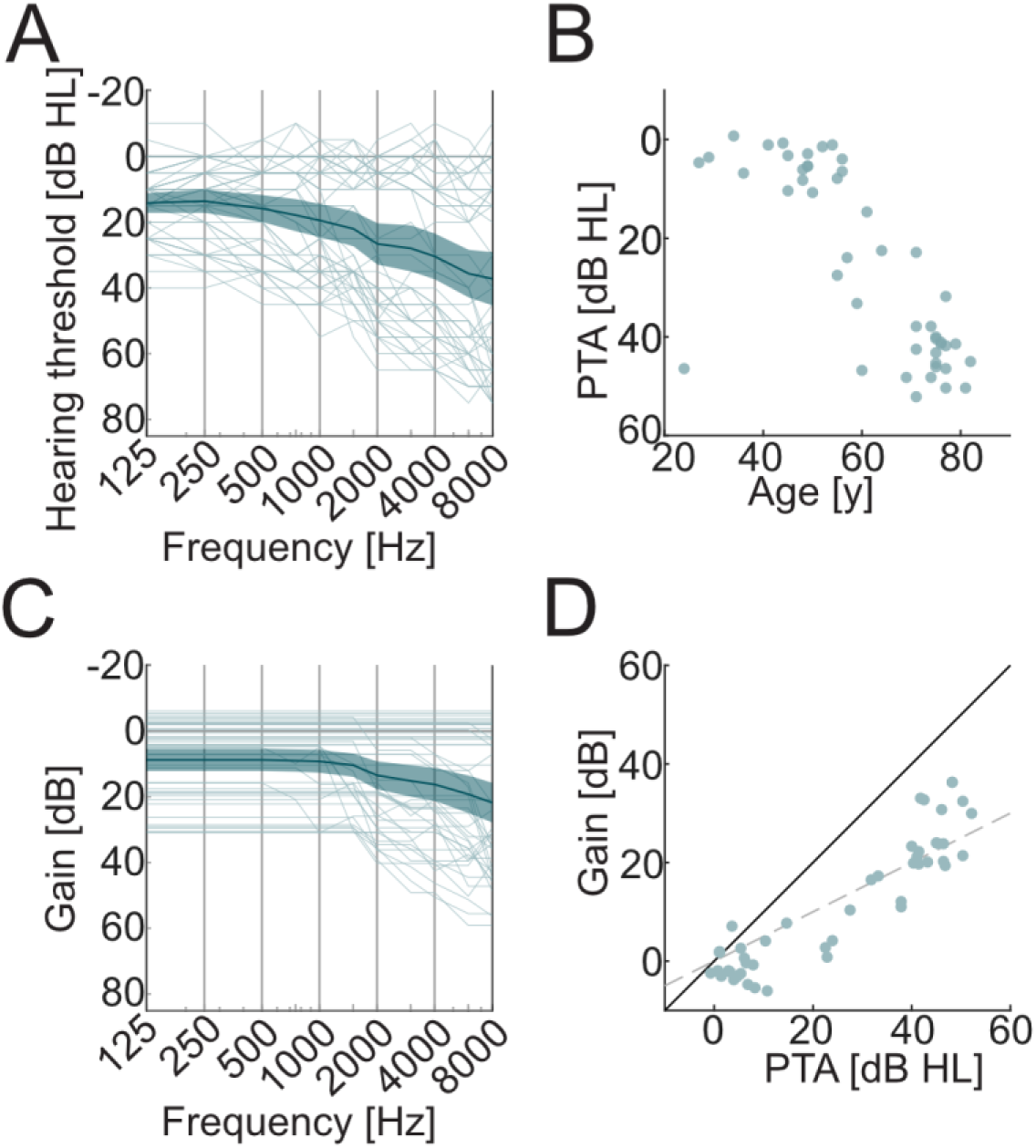
Hearing loss and compensation. Thin lines and dots depict single subjects. Thick lines and shaded areas depict mean and 95% confidence interval. **A)** Pure-tone hearing thresholds (air conduction) of the frequency-wise better ear shows heterogeneity of audiograms. **B)** Scatter plot of age and pure tone average (PTA) depicts collinearity. **C)** The depicted individual linear gain was applied to compensate for elevated hearing thresholds. First, frequency-dependent gain was applied to equalize for hearing thresholds. Second, an adaptive procedure was used to estimate the stimulus specific hearing threshold after equalization. **D)** Scatter plot of PTA vs. gain (average at PTA frequencies) illustrates that the resulting gain reached approximately half of the PTA (dotted line).

Next, a stimulus-specific hearing threshold was estimated to derive an individual broad-band amplification to be applied to the equalized stimuli. We estimated this by presenting the participants with a total of eight 30-second examples of the target stimuli (two from each talker), to which the gain was applied as derived in step one. The presentation started below hearing level and the level was increased at a rate of 2 dB per second. We asked the participants to press the space bar as soon as they could hear the slightest whisper. The average amplification at which they pressed the button was calculated and applied as an additional broad-band amplification, which will be referred to as individual hearing level (IHL). The overall resulting gain is depicted in Figure 2C. The comparison of the PTA and the applied gain suggests that the resulting amplification approximately fits the one-half-gain rule (Fig. 2D; Lybarger, 1944). As a sanity check, the adaptive procedure was repeated but this time we asked participants to press the space bar as soon as they could understand 95% of the words, which will be referred to as comprehension level (CL). This was to make sure that the presentation level was above the individual CL, or in other words, that participants could understand the target stimuli. We found that the individual difference between CL and IHL varied between 7 and 29 dB. For all participants, sounds were presented at 40 dB above IHL.

### Procedure and task

Experiments were conducted at Eriksholm Research Centre (Snekkersten, Denmark). Before the experiment, potential participants were asked to carefully read a document about the procedure and the related risks. At the beginning of their visit, participants gave written informed consent. The whole visit took between 2.5 and 3 hours.

A pure-tone audiogram was conducted at the beginning of the test session. Based on the audiogram, the individual gain was calculated (see Hearing loss compensation and adaptation). Then, the adaptive procedure for finding the IHL was conducted. After that, the participants were fitted with an EEG cap and two additional electrodes were secured to their mastoids. The electrodes were prepared with electrolyte gel. To ensure proper connection between electrodes and scalp (indicated by an offset-voltage > 50 mV or noisy signals), EEG signals and offset-voltages were checked, and additional gel was applied if necessary. The eye tracker was adjusted (such that it reliably captured both eyes) and calibrated using the five-dot calibration procedure in Tobii Eye Tracker Manager.

The main experiment started with six training trials (see above), initiated by the participant pressing “Start” on the computer keyboard. After each trial, participants were prompted with two statements about the content of the newsclips. They were asked to indicate whether each statement was true using the computer keyboard buttons labelled with “Ja” (Yes) or “Nej” (No). The performance in this task will be referred to as ***target comprehension***. After the response to the second question, the next trial started automatically. After training, participants were asked if the loudness was at a comfortable level, which all participants confirmed. If needed, EEG electrodes or the eye tracker were re-adjusted. Next, participants underwent four blocks of testing, each consisting of 10 one-minute trials. Between each block, participants were offered a break. On average, the break lasted for around 3 minutes.

After the four experimental blocks, it was revealed that there was an additional surprise recognition task of about 10 minutes. Participants were presented with 200 distractor sounds. Half of the 200 sounds were presented during the main experiment, whereas the other half were not. After each sound, a question mark appeared on the screen and participants were asked to indicate (using the “Yes” and “No” keys) whether they recognized hearing that sound in the main experiment. After they pressed a key, the next sound was presented between 1 and 2 seconds later (jittered interval). Subjects performed barely above chance in the surprise recognition task (Mean: 52.3%, SD: 4.2%), such that we did not consider the data for further analysis.

### Pupillometry

Task-evoked pupil dilation has been shown to depend on cognitive effort (Hess & Polt, 1964; Kahneman & Beatty, 1966), most likely due to phasic modulation of arousal, as controlled by noradrenergic activity in the locus coeruleus (Aston-Jones & Cohen, 2005; Joshi et al., 2016; Reimer et al., 2016). Effort and attentional control have been closely linked in the literature (Kahneman, 1973; for review: van der Wel & van Steenbergen, 2018). In animals noradrenaline decreases signal-to-noise ratio in firing patterns (Foote et al., 1975) and sharpens frequency tuning (Manunta & Edeline, 2004) of neurons in auditory cortex. In humans, pupil dilation during comprehension of sentences has been shown to correlate with task demands (for review: Kuchinsky & DeRoy Milvae, 2024; M. B. Winn et al., 2018). For example, lower signal-to-noise ratio (Kramer et al., 1997; Ohlenforst et al., 2017, 2018), spectral degradation (M. B. Winn et al., 2015), and higher semantic complexity (Kadem et al., 2020; M. B. Winn, 2016) lead to a larger pupil responses. Hence, pupil responses indicate the cognitive effort related to listening, or brief listening effort (Pichora-Fuller et al., 2016). Conversely, task-relevance has a strong effect on the pupil response to sentences (Pielage et al., 2021): while task-relevant sentences lead to a typical pupil dilation, task-irrelevant sentences only lead to small, barely above zero dilations. This indicates that the pupil response is mainly driven by cognitive processes rather than sensory input alone. Pupil responses to task-irrelevant sounds other than speech have been documented during visual tasks (Cronin et al., 2023; Hebisch et al., 2024; Petersen et al., 2017; Tona et al., 2016). Whether pupil responses to task-irrelevant sounds reflect distraction during an auditory task has not been documented to the same extent: There have been attempts to investigate how the pupil response reflects salience, but only in response to stimuli that were explicitly task relevant (Huang & Elhilali, 2017; Liao et al., 2016; Zhao et al., 2019). While stimulus-driven attention is initially automatic and may affect phasic arousal (Rajkowski et al., 1994), a reorientation of voluntary attention back to the target requires cognitive control, hence additional effort. The more salient a distractor is, the more effort will be needed to reorient attention back to the target, hence, the greater the pupil response should be.

We recorded pupillometric data at a sampling rate of 1200 Hz. For each trial, the raw pupillometric data consisted of vectors, one for each eye, indexing pupil size in mm as well as missing values (Not a Number, NaN) at timepoints where the pupil could not be detected. Missing values could occur due to blinks, but also other noise in the image processing during the pupil detection. It is common practice to remove samples before and after consecutive missing values to remove artifacts related to blinks (M. B. Winn et al., 2018). We applied this rule only to consecutive missing values longer than 20 ms (i.e., 24 samples). For these instances, we set the samples within the time range of 35 ms before (i.e., 40 samples) and 100 ms after (i.e., 120 samples) to missing values (NaNs). After this procedure, we still found spikes in the data such that we applied another rule based on the magnitude of the difference between successive samples. For each trial, the standard deviation (SD) of the sample-to-sample difference was calculated and successive samples with an absolute difference that exceeded two SDs were both set to NaN. All NaNs were linearly interpolated and the pupillometric data were low-pass filtered using a one-second-wide Hamming window.

Inspection of average pupil time courses across the trial revealed a strong inter-subject variability in pupil size (Fig. 3A) and a negative-going drift across the trial (Fig. 3B). To control for potential confounds, such as a subject with generally larger baseline pupil size also showing larger responses to distractors, two covariates were extracted from pupil size (Fig. 1C): *trial baseline pupil size* refers to the mean pupil size in the trial baseline period between −3 to 0 seconds relative to target speech onset. *Trial pupil size drift* refers to the mean pupil size difference from trial baseline pupil size during the trial between 0 and 60 seconds relative to target onset. To minimize the impact of drift on the extraction of pupil responses, trial-specific pupil time courses, within subject, were mean-corrected by subtracting the mean time course calculated across all trials for that subject.

**Figure 3:**
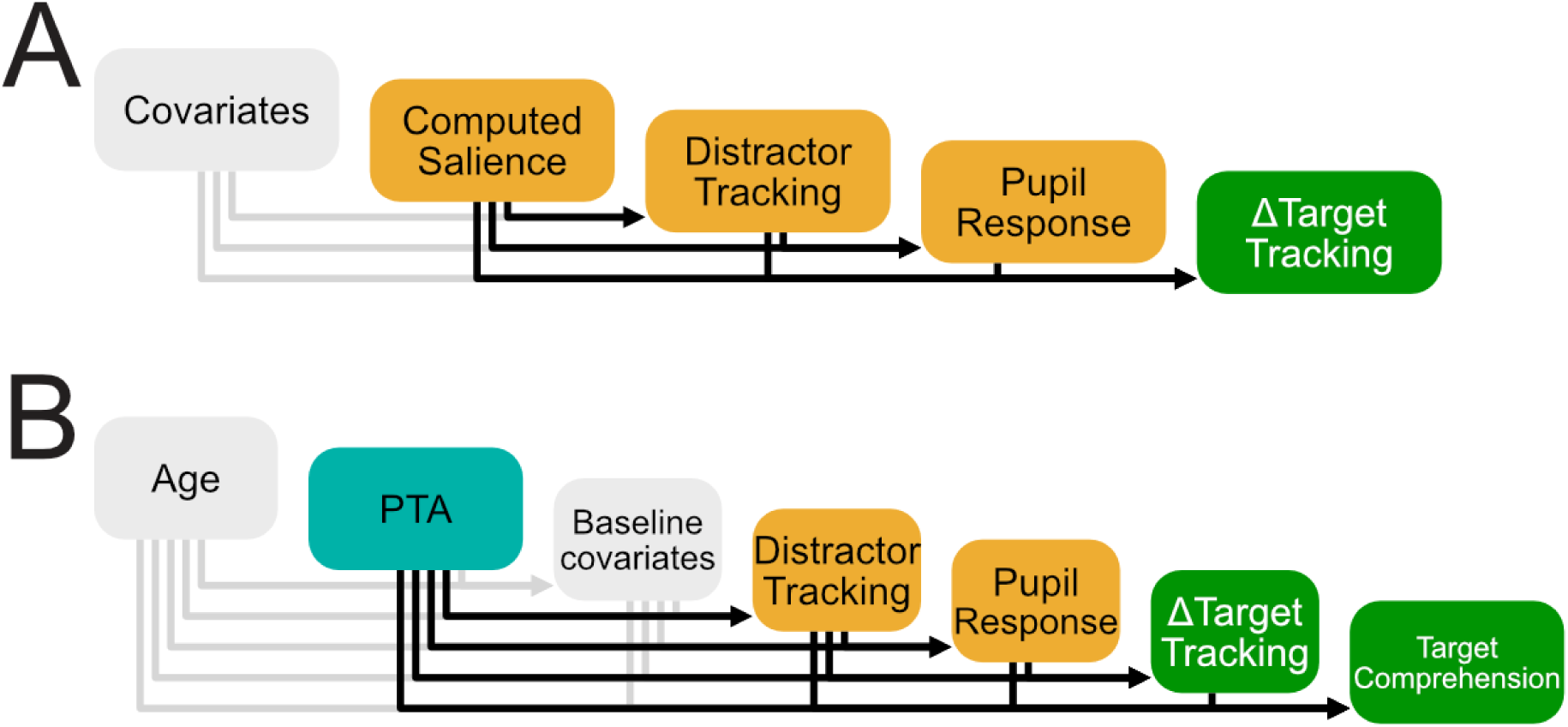
Hierarchical Statistical Modelling: **A)** Within-subject statistical modelling estimated the effect of stimulus features of single distractor sounds (computed salience) on single neurophysiological responses (distractor tracking, pupil response, Δtarget tracking) as well as their interdependence, while controlling for baseline measures (covariates, see figure 3-1A). **B)** Between-subject statistical modelling estimated the effect of hearing loss on subjects’ average neurophysiological responses, and their common effect on target comprehension, while controlling for covariates (age, average baseline covariates, see figure 3-1B).

To extract the pupil size time course around the distractor sounds, the preprocessed pupil size data were epoched between −1 and 5 seconds relative to distractor onset. The mean of the pupil size in the baseline interval between −1 and 0 seconds (***baseline pupil size***) was subtracted from the pupil size time course and stored as a covariate. The main measure of interest, the *pupil response* to distractors, was defined as the mean within the interval between 1 and 3 seconds. We only kept pupil responses in which less than 30% of the datapoints were missing before interpolation. Using this procedure, we obtained ***pupil responses*** to an average of 90.3% of the distractors. More than 80% of the pupil responses were available for 39 participants; between 50% and 80% were available for 7 participants; and only 21% were available for 1 participant. In sum, 11316 pupil responses were available for subsequent analysis.

### Electroencephalography

EEG data were recorded from 66 channels at a sampling rate of 8192 Hz. The preprocessing was mainly done with the FieldTrip-toolbox (version 20170321, Oostenveld et al., 2011), as well as custom-written code using MATLAB R2020b.

All the preprocessing was done on continuous data to avoid edge artifacts from filters. First, EEG data were re-referenced to the average of both mastoid electrodes. Second, the EEG data were resampled to 200 Hz. Next, a bidirectional band-stop filter (Butterworth, filter order: 6, cutoff frequencies: 47 & 53.2 Hz) and a bidirectional band-pass filter (FIR, cutoff frequencies: 1 Hz & 80 Hz, Hamming window, order: 600) were applied. The EEG data were epoched according to the markers sent at the beginning and the end of a trial. The EEG data were visually inspected, and EEG channels with obvious noise were identified, removed, and interpolated from neighboring channels. While 26 subjects had no bad channels at all, 20 subjects had between 1 and 3 bad channels and 1 subject had 5 bad channels.

Next, an independent component analysis (Makeig et al., 2004) was performed using the *runica* algorithm, as implemented in FieldTrip. The number of independent components (IC) was set to N_IC_ = N_C_ – N_BC_ – 1, where N_C_ is the number of EEG channels (i.e., 66) and N_BC_ is the number of bad channels. The IC were visually inspected both in the time and frequency domain, in addition to the topographic distribution of their weights. IC that could be clearly related to eyeblinks, eye movements, muscle artifacts, heartbeat, or single channel noise were removed from the EEG data. On average, 16.62 (SD: 6.35) IC were removed. To make sure that we have not accidentally removed any auditory components, we compared event-related potentials to noise and target onsets before and after cleaning of IC and did not observe a reduction in the ERP components. Finally, EEG data were low-pass filtered at 10 Hz using a bidirectional Butterworth filter (order: 16).

### Neural tracking of target speech and distractor sounds

Neural tracking can be extracted from electrophysiological signals such as electroencephalography (EEG). It quantifies the strength of the overall neural response to a continuous stimulus feature such as the envelope. In numerous studies, it has been shown that attended auditory stimuli are neurally tracked more strongly than ignored stimuli (Ding & Simon, 2012; Fiedler et al., 2017, 2019; Tune et al., 2021). These studies mainly investigated the effect of sustained voluntary attention on neural tracking, but not the effect of brief, stimulus-driven disruption of attention. Only recently, researchers have begun to investigate how events in the to-be-ignored stream, such as changes in background noise (Khalighinejad et al., 2019), or sudden brief sounds (Holtze et al., 2021), affect the neural tracking of to-be-attended speech, also with regards to their computed salience (Straetmans et al., 2021). Here, we ask if the computed salience of to-be-ignored sounds correlates with the neural tracking of these sounds, as well as with the attention-capture-related perturbation of neural tracking of target speech.

We calculated neural tracking of both distractor sounds and target speech to analyze how strongly the stimuli were represented in the EEG and how much this representation was modulated by attention. We followed the well-established approach of temporal response functions (TRF; Crosse et al., 2016) using the mTRF-toolbox version 2.3 (https://github.com/mickcrosse/mTRF-Toolbox). We used both a forward (encoding) model to illustrate TRFs together with their scalp distributions as well as backward (decoding) models for the estimation of scalp-wide neural tracking and statistical comparisons.

Both the forward and backward approach follow the assumption that a linear filter (i.e, TRF) can be estimated that, convolved with a predictor variable, describes some variance in a response variable, while the rest is noise:

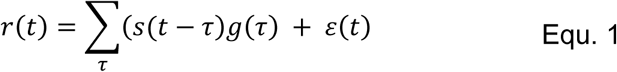

where *t* is time points, *s* is the predictor variable, *g* is the TRF, *τ* is time lags between the predictor and the response, *ε* is noise, and *r* is the response variable. To estimate the *g* that minimizes the mean-squared error, a matrix operation can be performed that resembles multivariate linear regression:

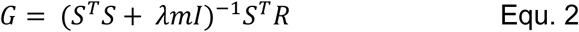

where *S* is a matrix containing the predictor variable(s) in columns (as well as its time-lagged replications based on *τ*), *T* denotes matrix transposition, *m* is the mean trace of the covariance matrix *S^T^S* (Biesmans et al., 2016)*, λ* is the regularization parameter, *I* is the identity matrix, *R* is the response variable and *G* is the TRF. For forward models, the predictor variable is a stimulus feature and the response variable is the EEG, whereas for backward models it is the other way around.

As a basis for the extraction of a stimulus feature, we used the stimulus sound files. We did not measure the stimuli at ear level, but since the loudspeakers have a fairly flat frequency response (deviation of ±2 dB between 58 Hz and 20 kHz, Genelec Oy, 2007) and the listening booth is acoustically treated, we did not expect substantial distortion. For target speech stimuli, we used the two concatenated 30-second newsclips as they were presented per trial. In case of the distractor sounds, we used the waveform that resulted from embedding the distractor sound into the noise as it was presented at the left or right channel with 5 seconds of noise before and after distractor onset to account for edge artifacts. Note that neural tracking of the distractor will only be calculated for the 1-second time window of the distractor (see below).

As stimulus feature, envelope onsets were extracted from the stimuli by following procedures as in Fiedler et al., 2019: First, a gammatone filter bank with 39 bands between 50 Hz and 15 kHz was used. Second, the absolute value of the analytic signal of each band was derived. Third, the same low-pass filter (same as for the EEG: Butterworth, cutoff: 10 Hz, order: 16) was used on each band. Fourth, the first derivative was taken in each band and negative values were set to zero (i.e., halfwave rectification). Finally, the resulting signals were summed across bands.

We calculated neural tracking of both target speech and distractor sounds separately, but in a similar way. We used a within-subject leave-one-out cross validation approach consisting of three steps: First, the TRFs (both forward and backward) were trained on the envelope onsets and EEG data from all but one trial between time lags (*τ*) of −150 and 550 ms (Equation. 2). To this end, envelope onsets and EEG were concatenated with 550 ms of zeros in between to avoid any leakage. For computational efficiency, in the case of backward models, the envelope onsets and EEG were resampled to 50 Hz. The regularization parameter *λ* was set to reduce edge- and sample-to-sample artifacts in the TRFs (Fiedler et al., 2019). While we set *λ* = 0.1 for the forward model, we set *λ* = 1 for the backward model due to the higher covariance among EEG channels, which requires more regularization. Second, the TRF was used to predict EEG (forward) or to reconstruct the envelope onsets (backward) of the left-out trial. Lastly, the Pearson-correlation coefficient between the predicted and the measured EEG signal (forward) and between the presented and reconstructed envelope (backward) of the left-out trial was calculated. This correlation coefficient depends directly on the strength of the neural representation of the envelope onsets in the EEG, and hence, will be referred to as neural tracking.

Regarding the distractor sounds, the goal was to calculate their neural tracking (*distractor tracking*) separately for each sound occurring within a trial within their presentation window of one second. The tracking of each distractor was calculated based on the prediction/reconstruction with the TRF trained on all sounds that occurred in all the other trials. To capture the full response in the EEG, we extended the cutout of envelope onsets and EEG from −150 ms relative to distractor onset to 550 ms relative to distractor offset, covering the range of time lags (as mentioned above). We observed a slight lateralization of distractor tracking (see Results): it was stronger at EEG electrodes contralateral to distractor presentation. To account for the lateralization, we mirrored the channel layout of the EEG data along the sagittal axis for distractors presented on the right, such that the topographies will be presented in ipsi- and contra-lateral fashion instead of left and right. To illustrate the extent of lateralization, we also calculated the difference in distractor tracking between ipsi- and contralateral electrode pairs.

Regarding the neural tracking of target speech (target tracking), we first estimated one-second-long windows of target tracking in steps of 0.1 seconds across whole trials for illustrative purposes. Second, to estimate the effects of distractor presentation on target tracking, we estimated target tracking for windows of two seconds pre-distractor onset (i.e., −2 to 0 sec) and two seconds post-distractor onset (i.e., 0 to 2 sec). The pre-distractor target tracking will be referred to as *baseline target tracking*, and the pre-to-post change in target tracking will be referred to as *Δtarget tracking*. After inspecting the windowed target tracking around distractor presentation, we observed artifacts that resulted from consistent event-related potentials in response to the distractors. To control these artifacts, the average ERP to distractor onsets of each subject was subtracted from their individual data before target tracking was estimated, which resulted in a smooth, artifact-free estimation of target tracking.

### Salience computation

We used an existing, behaviorally validated model for acoustic salience (Huang & Elhilali, 2017). In brief, this model was developed to predict human decisions about which of two concurrent streams, each presented to one ear, is capturing the listeners attention moment-to-moment. Psychoacoustic features were extracted from the streams and underwent a factor analysis. Unsurprisingly, loudness was found to be the strongest driver for salience, but other factors such as harmonicity also play a role. Here we rms-equalized the distractor sounds (see Stimulus material), such that their variability in loudness was minimized and other features dominated the variability in overall computed salience.

We submitted the distractor sounds embedded in the stationary noise to the salience model as provided by Huang & Elhilali (2017). We used the default parameters. The salience model returned a salience time course, which we averaged across the 1-second window in which the distractor was inserted. Hence, we ended up with one value of *computed salience* for each distractor sound. These computed salience values were transformed into z-scores.

### Statistical analyses

We tested if predictors (such as computed salience) affected our response variables (such as neurophysiological responses or behavior) in the manner we hypothesized. Additionally, we controlled for several covariates, as described below.

We used a hierarchical series of (generalized) linear mixed models to explain variance of dependent variables (i.e., responses) by independent variables (i.e., predictors). Residuals of earlier models were used as predictors for subsequent models. For example, we first tested if computed salience predicted distractor tracking. After we found a significantly positive estimate for modelled salience on distractor tracking, we regressed it out from distractor tracking and used the residuals of distractor tracking as a predictor for pupil response. This way, a common modulation by computed salience of both distractor tracking and pupil response would not spuriously lead to a positive estimate of distractor tracking on distractor-evoked MPD. Only if distractor tracking explained additional variance beyond the variance that computed salience explained directly, would a significant relationship be revealed.

Besides the variables of interest (Fig. 3), we used several covariates, such as trial number and baseline of the neurophysiological responses (Fig. 3-1). Each response was modelled in a stepwise (generalized) linear mixed modelling approach with random and fixed effects. We started with a basic model and added predictors one at a time (i.e., increased model). At each step, a likelihood-ratio test between the increased model and the current model was performed to determine if a predictor significantly improved the model fit (p_lrt_ < 0.05 and improved r-squared). If this was the case, the predictor was added to the current model. When a predictor was added, all previously included predictors were removed one at a time (i.e., reduced model). If their removal did not significantly degrade the model fit (according to a likelihood-ratio test between the current and the reduced model), the predictor was removed from the model. That way we ensured that we accounted for the collinearity of some predictors such as age and PTA.

We ran this stepwise procedure on two levels: 1) the within-subject level: here we modelled the neurophysiological and behavioral response to single distractor sounds (Fig. 3A, Fig. 3-1A). 2) the between-subject level, where predictors and response variables only varied between subjects, such as age, means of the neurophysiological and behavioral measures (Fig. 3B, Fig. 3-1B). This way, we ensured that we could distinguish between interpretations on the within-subject level, such as “A relatively strong response to a unique distractor indicates that a subject was more distracted by this particular sound compared to another sound where a weaker response was found” and the between-subject level such as “A subject with a strong average response to distractor sounds was more distractable than another subject showing a weaker response”.

Since age (and, to a lesser extent, PTA) has been previously shown to affect both pupil size (B. Winn et al., 1994) and neural tracking (Karunathilake et al., 2023; Kulasingham et al., 2020; Zan et al., 2020), there may be inter-individual differences in the dynamic range of these measures. We accounted for these potential differences on both the within- and the between-subject level: On the within-subject level, we z-scored each subject’s measures relative to their own means and standard deviations across all trials. On the between-subject level, we controlled for potential differences in the overall level and dynamic range of pupil size and neural tracking by including the individual standard deviation of trial baseline pupil size (*trial baseline pupil size dynamics*) and baseline target tracking (*baseline target tracking dynamics*) as covariates in the hierarchical models. Note that this way, age and PTA were first tested on the baseline covariates as well as the measures of interest, and observed effects were regressed out, such that potential confounds were controlled for. For example, we indeed found that older participants showed higher baseline target tracking (see “Dependencies among covariates”). After this effect was regressed out, we found that higher baseline target tracking indicated better target comprehension. This can be interpreted as demonstrating that greater target tracking (for a given age) leads to better target comprehension.

To summarize, we modelled neurophysiological models on both the within- and between subject level under consideration of several covariates. On the within-subject level, we followed the order as depicted in (Fig. 3A, Fig. 3-1A): trial number, baseline pupil size, baseline target tracking, computed salience, distractor tracking, pupil response, and Δtarget tracking. On the between-subject level, we followed the order as depicted in (Fig. 3B, Fig. 3-1B): Age, PTA, trial baseline pupil size, trial baseline pupil size dynamics, trial pupil size drift, baseline target tracking, baseline target tracking dynamics, distractor tracking, pupil response, Δtarget tracking and target comprehension.

## Results

We asked human subjects to listen to continuous speech while distractor sounds of varying computed salience were presented at unpredictable times and locations. We statistically modelled both the neurophysiological and behavioral responses to the distractors and target speech to test if they can be interpreted as a proxy of distraction. Furthermore, we tested if distractibility increases with hearing loss. Before reporting the main results, the dependencies among the covariates will be described.

### Dependencies among covariates

To control potential confounds, we first investigated the effects of trial number, age, and PTA on baseline covariates such as baseline pupil size and baseline target tracking.

An illustration of dependencies among the covariates (as well as the main measures of interest) can be found in figure 4-1. Slopes are depicted in figure 6-1 (pupil size) and 7-1 (target tracking). Within subjects (Fig. 4-1A), baseline pupil size decreased over the course of the experiment: trial number negatively predicted baseline pupil size (χ^2^(1) = 2483, p_lrt_ = 2.2*10^-308^, ß = −.446, SE = 8.46 ^-3^, t(11314) = −52.7, p = 4.94*10^-324^, partial r^2^ = .197). Between subjects (Fig. 4-1B), older participants showed both smaller and less variable pupil size compared to younger ones: Age negatively predicted both trial baseline pupil size (χ^2^(1) = 21.1, p_lrt_ = 4.28*10^-6^, ß = −3.61*10^-2^, SE = 6.98*10^-3^, t(45) = −5.17, p = 5.28*10^-6^, partial r^2^ = .362) and trial baseline pupil size dynamics (χ^2^(1) = 11.6, p_lrt_ = 6.53*10^-4^, ß = −1.42*10^-3^, SE = .39*10^-3^, t(44) = −3.63, p = 7.2*10^-4^, partial r^2^ = .219). Subjects with higher PTA showed a smaller negative pupil size drift over the course of the trial: PTA negatively predicted trial pupil size drift (χ^2^(1) = 5.52, p_lrt_ = 0.0188, ß = 4.5*10^-3^, SE = 1.16*10^-3^, t(44) = 3.879, p = 3.47*10^-4^, partial r^2^ = 0.243). Additionally, subjects with higher trial baseline pupil size dynamics showed a larger negative trial pupil size drift as indicated by a negative estimate (χ^2^(1) = 11.32, p_lrt_ = 7.6610^-4^, ß = −1.86, SE = .520, t(44) = 3.58, p = 8.57*10^-4^, partial r^2^ = 0.214). Older subjects showed higher baseline target tracking as indicated by a positive estimate (χ^2^(1) = 10.52, p_lrt_ = 0.012, ß = 8.63*10^-4^, SE = 2.51*10^-4^, t(45) = 3.43, p = 1.30*10^-3^, partial r^2^ = 0.201). Subjects with higher PTA showed higher variability in baseline target tracking: PTA positively predicted baseline target tracking dynamics (χ^2^(1) = 5.72, p_lrt_ = 0.0167, ß = 1.16*10^-4^, SE = 4.72*10^-5^, t(45) = 2.47, p = 0.0175, partial r^2^ = 0.115).

**Figure 4:**
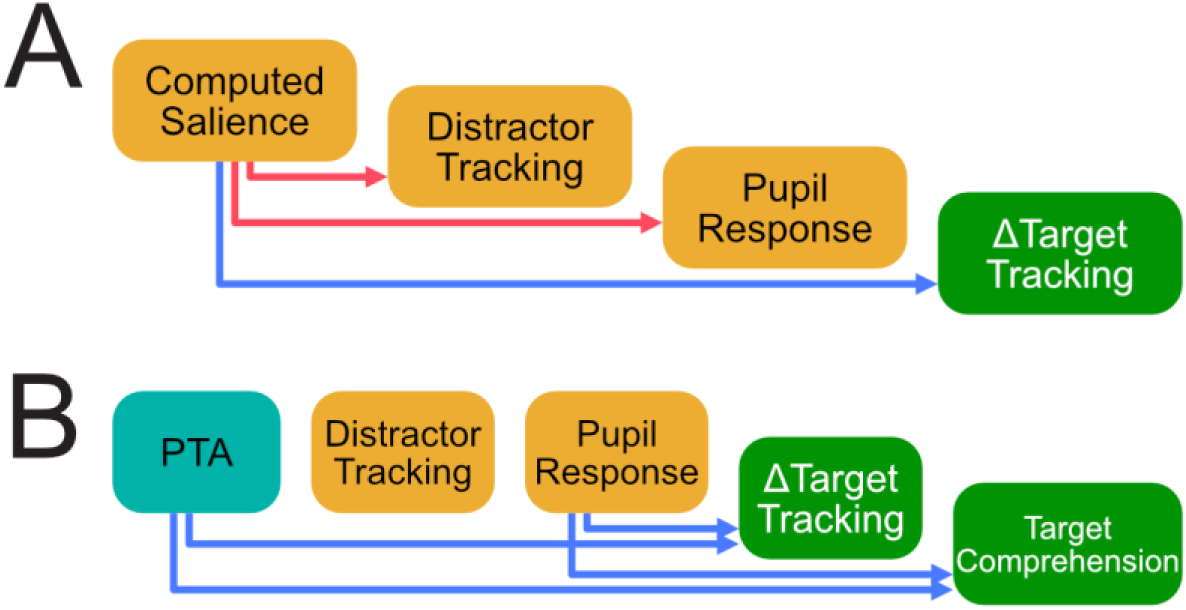
Results: Effects among measures of interest. Blue arrows indicate negative relationships, red arrows indicate positive relationships. **A)** Within-subject model confirm expected effects: More salient sounds lead to stronger distractor tracking and pupil response, while a more negative drop in target tracking (Δtarget tracking) can be observed. **B)** Between-subject model reveals that mainly pupil response indicates distractibility: Subjects with larger pupil response showed a stronger drop in target tracking (Δtarget tracking) accompanied by worse target comprehension. Models were controlled for several covariates and their relationships are depicted in figure 4-1.

In sum, we found the expected effects of age and PTA on the baseline measures: Higher age leads to smaller, less variable pupil size, whereas higher age (and PTA) lead to stronger (and more variable) target tracking. Any significant predictors among the covariates were regressed out and the residuals were used for further prediction of the main measures of interest (see methods for more details).

### Distractor tracking

Descriptively, we found consistent temporal response functions to distractors with a strong positive component at around 250 ms (Fig. 5A). Above-zero distractor tracking was observed in all participants (Fig. 5B), which was confirmed by a significant intercept to be above zero in the between-subject model (ß = .150, SE = 4.73*10^-3^, t(46) = 31.6, p = 7.633*10^-33^). Strongest distractor tracking was observed at frontocentral channels with a slight lateralization indicating stronger tracking at channels contralateral to distractor presentation, which has been controlled for in the estimation of distractor tracking (see methods).

**Figure 5:**
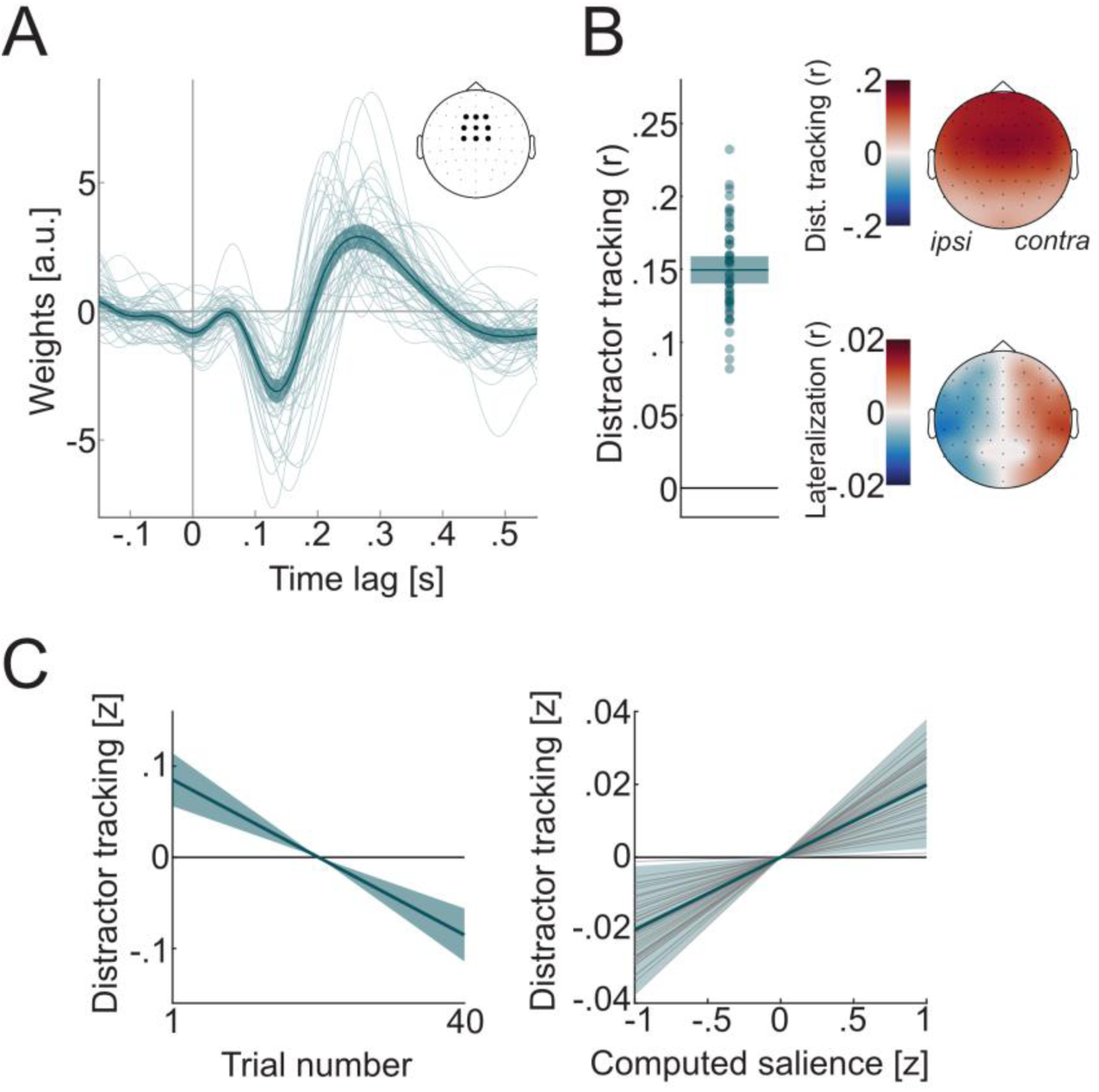
Neural tracking of distractor sounds (distractor tracking). Thick lines depict the mean across subjects (A,B) or slope estimates of fixed effects (C). Shaded areas depict 95% confidence intervals of the mean or slope estimates. Thin lines and dots depict single subject means (or random slopes if stated). **A)** Temporal response functions in response to distractors derived from forward modelling averaged across frontocentral channels as indicated by topography. **B)** Distractor tracking derived from backward modelling and topographies derived from forward modelling depicting the scalp distribution and lateralization. **C)** Within-subject predictors of trial number (centered) and computed salience (z-scored). Thin grey lines depict individual random slopes (which were only estimated for computed salience for illustration).

Within subjects (Fig. 5C), the model included the covariate trial number (χ^2^(1) = 32.73, p_lrt_ = 1.06*10^-8^) and computed salience χ^2^(1) = 32.73, p_lrt_ = 1.06*10^-8^). Distractor tracking decreased over the course of the experiment: Trial number negatively predicted distractor tracking (ß = −.51, SE = 8.91*10^-3^, t(12531) = −5.73, p = 1.01*10^-8^, partial r^2^ = .0026). As hypothesized, distractor tracking was stronger for sounds with higher computed salience (ß = 1.99*10^-2^, SE = .891*10^-2^, t(12531) = 2.23, p = .0256, partial η^2^ = .0004). The final model explained only a small amount of the variance (R^2^_adj_ =.0028).

On the between-subject level, none of the predictors were significant, such that none of the inter-subject variance could be explained by the model (R^2^_adj_ = 0).

In sum, distractor tracking depended positively on computed salience. The only significant within-subjects covariate was trial number, which may indicate that distraction decreased over the course of the experiment. Surprisingly, between subjects, neither increased age nor PTA led to increased distractor tracking. Furthermore, there was also no positive relationship between baseline target tracking and distractor tracking, which may be expected given that some subjects generally show stronger neural tracking than others.

### Pupil response

Descriptively, we observed variability in trial baseline pupil size across subjects ranging from 2 to 6 mm (Fig. 6A). Baselined trial time courses revealed a negative drift (trial pupil size drift) of pupil size (Fig. 6B), which was confirmed to be below zero by a significant intercept in the between-subject model (ß = −.156, SE = 0.0219, t(44) = −7.11, p = 7.87*10^-9^). We found consistent pupil responses to distractors peaking at around 1.5 seconds (Fig. 6C). Pupil response was confirmed to be above zero by a significant intercept in the between-subject model (ß = .043, SE = 2.72*10^-3^, t(45) = 15.8, p = 5.59*10^-20^).

**Figure 6:**
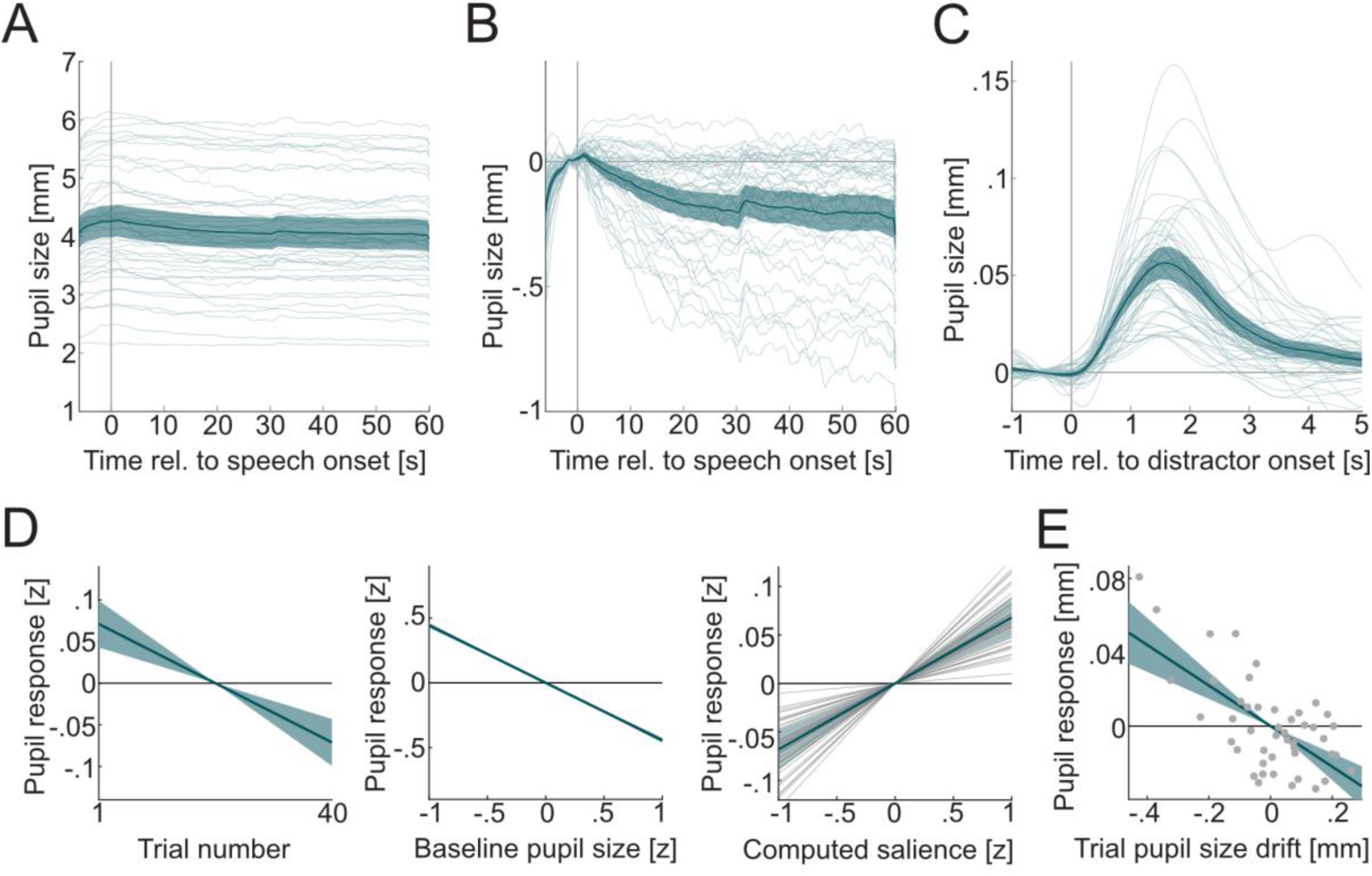
Pupil size and pupil response. Thick lines depict the mean across subjects (A-C) or slope estimates of fixed effects (D,E). Thin lines and dots depict single subject means (or random slopes if stated). **A)** Pupil size across trial. **B)** Baselined pupil size across trial with negative drift (trial pupil size drift). Pupil response at around 30 seconds indicates change of target talker. **C)** Pupil response curve to distractor sounds. Mean pupil dilation (i.e., pupil response) was calculated between 1 and 3 seconds. **D)** Within-subject predictors of pupil response (all z-scored but trial number): Trial number (centered), baseline pupil size (confidence band very small), computed salience (thin grey lines depict random slopes). **E)** Between-subjects predictors of pupil response (centered): Trial pupil size drift (centered). Effects on pupil size covariates are depicted in figure 6-1.

Within subjects (Fig. 6D), the model for pupil response included trial number (χ^2^(1) = 20.2, p_lrt_ = 7.06*10^-6^), baseline pupil size (χ^2^(1) = 1945, p_lrt_ < 2.23*10^-308^), and computed salience (χ^2^(1) = 63.02, p_lrt_ = 0.0141). Pupil response decreased over the course of the experiment: trial number negatively predicted pupil response (ß = −4.27*10^-2^, SE = .863*10^-2^, t(11312) = −4.94, p = 7.82*10^-7^, partial r^2^ = .0024). Relatively large pupils just before the distractor sound lead to smaller pupil response (and vice versa): baseline pupil size negatively predicted pupil response (ß = −.444, SE = 9.59*10^-3^, t(11312) = −46.29, p < 2.23*10^-308^, partial η^2^ = .150). As hypothesized, pupil response was larger for sounds with higher computed salience: computed salience positively predicted pupil response (ß = 6.78*10^-2^, SE = 1.07*10^-2^, t(11312) = 6.33, p = 2.56*10^-10^, partial r^2^ = .0061). The final model explained a considerable amount of within-subject variance (R^2^_adj_ = .165).

Between subjects (Fig. 6E), the model for pupil response only included trial pupil size drift (χ^2^(1) = 27.1, p_lrt_ = 1.94*10^-7^). Subjects with a more negative trial pupil size drift showed larger pupil response as indicated by a negative estimate (Estimate: ß = −0.11, SE = 0.018, t(44) = −6.05, p = 2.6*10^-7^, partial r^2^ = 0.438). The final model explained a considerable amount of the between-subject variance (R^2^_adj_ = .438).

In sum, pupil response depended positively on computed salience. However, within-subjects covariates such as time on task (here: trial number) and baseline pupil size should be considered as well in the modelling of a single pupil response. The strong dependency of the pupil response on baseline pupil size indicates a regression to the mean effect: A relatively high baseline pupil size is likely followed by a decrease in pupil size and vice versa. The fact that trial pupil size drift was considered as a covariate in the between-subject level may indicate that there is some ceiling effect: Subjects who exhibit a greater negative deviation from pupil baseline over the course of the trial have more head room for a distractor-related pupil response.

### Target tracking

Descriptively, we found consistent temporal response functions to target speech (Fig. 7A). We found consistent target tracking that was constant across the trial and driven by fronto-central channels (Fig. 7B). Baseline target tracking was confirmed to be above zero by a significant intercept in the between-subject model (ß = 0.104, SE = 3.96*10*^-2^, t(45) = 26.2, p = 7.27*10^-29^). More importantly, we found consistent drops in target tracking post distractor onset (Fig. 7C), which resulted in Δtarget tracking below zero as confirmed by significant intercept in the between-subject model (ß = −.0315, SE = 1.93*10^-3^, t(45) = −16.3, p = 4.85*10^-20^).

**Figure 7:**
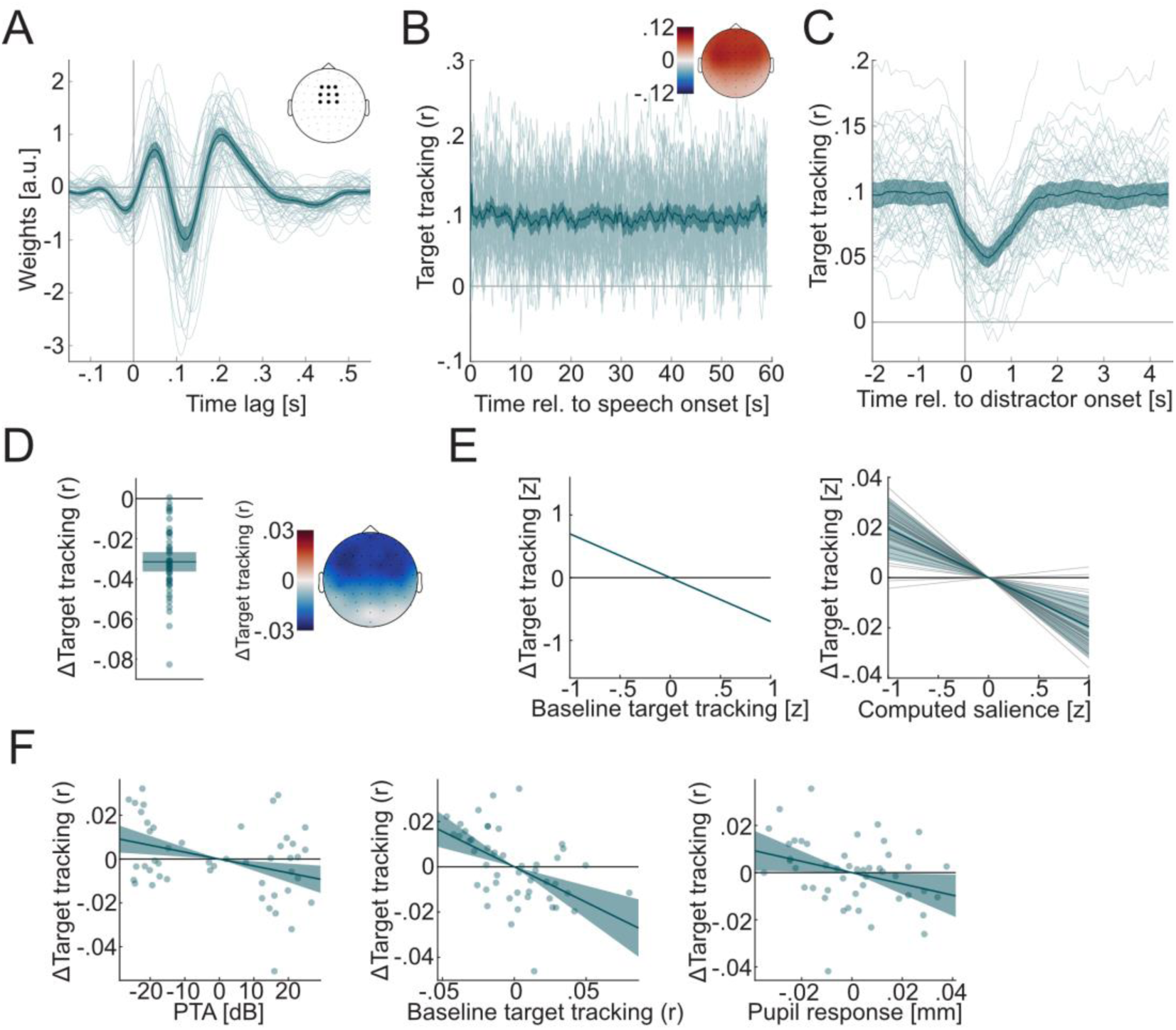
Target tracking. Thick lines depict the mean across subjects (A-D) or slope estimates of fixed effects (E,F). Thin lines and dots depict single subject means (or random slopes if stated). **A)** Temporal response functions in response to distractors derived from forward models averaged across frontocentral channels as indicated by topography. **B)** Windowed target tracking across trial derived from backward models (window size 1 sec, step 0.1 sec) and topography of target tracking derived from forward models. **C)** Distractor-related drop in target tracking illustrated by windowed target tracking. **D)** Δtarget tracking was estimated from backward models based on the difference between target tracking within windows 2 seconds before vs. 2 seconds after distractor onset. Topography of Δtarget tracking derived from forward models. **E)** Within-subject predictors of Δtarget tracking (all z-scored): baseline target tracking, computed salience (thin grey lines depict random slopes). **F)** Between-subject predictors of Δtarget tracking (all centered): PTA, baseline target tracking, pupil response. Effects on target tracking covariates are depicted in figure 7-1.

Within-subjects (Fig. 7E), the model for Δtarget tracking included baseline target tracking (χ^2^(1) = 8401, p_lrt_ < 2.23*10^-308^) and computed salience (χ^2^(1) = 9.40, p_lrt_ = 0.0022). Relatively strong target tracking just before the distractor sound led to a larger drop in target tracking: baseline target tracking negatively predicted Δtarget tracking (ß = −.699, SE = 6.39*10^-3^, t(12531) = −109.43, p < 2.23*10^-308^, partial r^2^ = .499). As hypothesized, the drop in target tracking was larger for sounds with greater computed salience: computed salience negatively predicted Δtarget tracking (ß = −1.96*10^-2^, SE = .638*10^-2^, t(12531) = 3.01, p = 2.17*10^-3^, partial r^2^ = 7.5010^-4^). The final model explained a considerable amount of variability in Δtarget tracking (R^2^_adj_ = .489), which could be mainly attributed to baseline target tracking.

Between subjects (Fig. 7F), the model included PTA (χ^2^(1) = 5.27, p_lrt_ = 0.0218), baseline target tracking (χ^2^(1) = 12.25, p_lrt_ = 4.65*10^-4^) and pupil response (χ^2^(1) = 4.66, p_lrt_ = .0309). Subjects with higher PTA showed a stronger drop in target tracking: PTA negatively predicted Δtarget tracking (ß = − 3.15*10^-4^, SE = 1.06*10^-4^, t(43) = −2.99, p = 4.63*10^-3^, partial r^2^ = .166). Subjects with higher baseline target tracking showed a larger drop in target tracking: baseline target tracking negatively predicted Δtarget tracking (ß = −.313, SE = .0728, t(43) = −4.30, p = 9.53*10^-5^, partial r^2^ = .292). Subjects with larger distractor-evoked MPD showed a larger drop in neural target tracking: distractor-evoked MPD negatively predicted Δtarget tracking with a negative estimate (ß = −.239, SE = .108, t(43) = −2.21, p = .0323, partial η^2^ = .103). The final model explained a moderate amount of the between-subject variance in Δtarget tracking (R^2^_adj_ = .333).

In sum, target tracking within subjects dropped when a distractor was presented, and the drop was larger for distractors with higher computed salience. The strong dependence of the drop on baseline target tracking suggests regression to the mean effects: a relatively high baseline in target tracking is likely followed by a decrease, and conversely, a low baseline is followed by an increase in target tracking. On the between-subject level, subjects with higher PTA and higher baseline target tracking also showed a larger drop in target tracking (Δtarget tracking). This may indicate a larger effect of distraction, but may also simply be an artefact: stronger baseline target tracking affords a larger drop.

### Target comprehension

The percentage of correctly answered comprehension questions was significantly above chance (> 59%) for each participant, which indicates everyone was able to follow the task instructions (Fig. 8A). This was also confirmed by a significantly positive intercept (ß = 1.82, SE = .0484, t(43) = 37.6, p = 1.56*10^-34^), which corresponds to a mean comprehension of 86.1 %.

**Figure 8:**
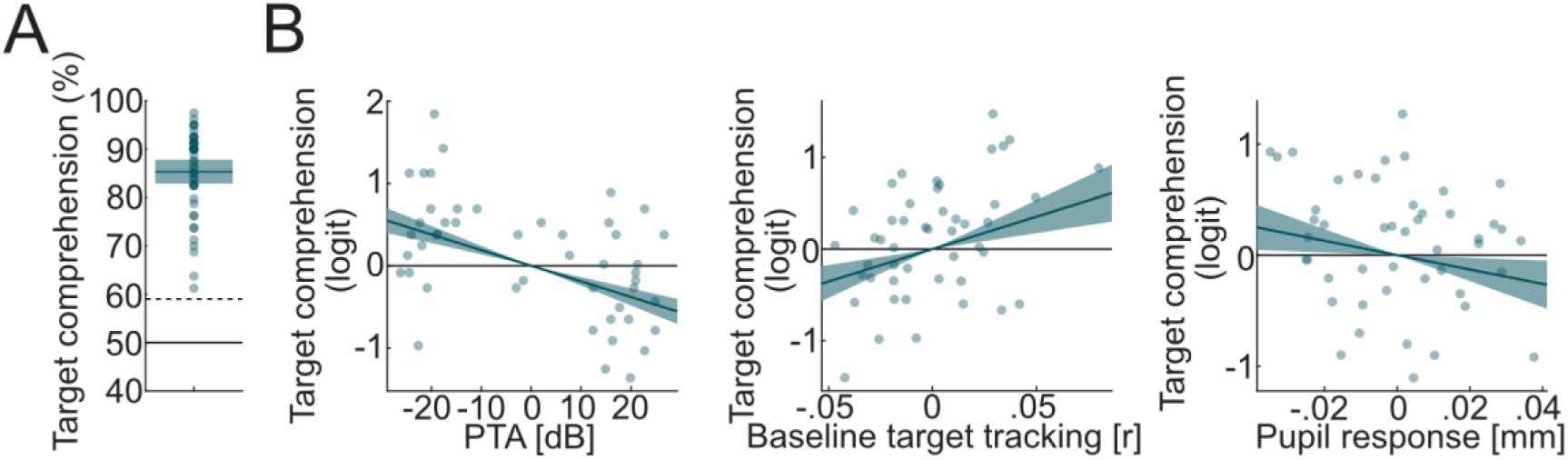
Target comprehension. Thick lines depict the mean across subjects (A) or slope estimates of fixed effects (B). Shaded areas depict 95% confidence intervals of the mean. Dots depict single subject means. **A)** Target comprehension as the percentage of correctly answered comprehension questions. Dotted line indicates significant performance above chance. **B)** Between-subject predictors of logit-transformed target comprehension (all centered): PTA, baseline target tracking, pupil response.

Between subjects (Fig. 8B), the model included PTA (χ^2^(1) = 13.7, p_lrt_ = 2.19*10^-4^), baseline target tracking (χ^2^(1) = 20.0, p_lrt_ = 4.64*10^-6^), and pupil response (χ^2^(1) = 5.73, p_lrt_ = .0167). Participants with higher PTA showed worse target comprehension (ß = −.0190, SE = 2.59*10^-3^, t(43) = −7.34, p = 4.17*10^-9^). Participants with higher baseline target tracking showed better target comprehension: baseline target tracking positively predicted target comprehension (ß = 7.07, SE = 1.78, t(43) = 3.96, p = 2.74*10^-4^). Crucially, participants with a larger pupil response showed worse target comprehension: pupil response negatively predicted target comprehension (ß = −6.47, SE = 2.61, t(43) = −2.48, p = 0.0169). The final model explained a considerable amount of between-subject variance in target comprehension (R^2^_adj_ = .290).

In sum, we found that higher PTA, higher pupil response, and lower baseline target tracking (for a given age) lead to worse target comprehension.

## Discussion

We tested whether neurophysiological responses to irrelevant sounds (distractors) indicate the degree to which attentional focus on continuous target speech is captured by such distractors, in a heterogenous group in terms of age and hearing loss profiles.

We controlled for the effects of age and hearing loss on baseline measures and responses. As expected, older subjects showed smaller, less variable pupils (B. Winn et al., 1994) and older subjects also showed stronger and more variable target tracking (Karunathilake et al., 2023; Kulasingham et al., 2020; Zan et al., 2020). Interestingly, after controlling for age we found that higher baseline target tracking leads to better target comprehension (Lesenfants et al., 2019; Vanthornhout et al., 2018), which may be related to sustained attention to the audiobook. The substantial within-subjects variability (due to trial number and baseline values) and between-subjects variability (due to age and PTA) had to be explicitly modelled before the predicted effects were apparent.

Regarding our hypothesis, we found that both distractor tracking and pupil response, as well as a drop in target tracking (Δtarget tracking) could be predicted from salience of the distractors. While subjects with stronger pupil responses also showed stronger drop in target tracking, only pupil response predicted target comprehension: subjects with stronger pupil response showed weaker target comprehension. While (compensated) hearing loss did not affect the distraction-related responses, a hearing-loss-related decrease in target speech comprehension was observed.

### Neural tracking and pupil response are salience-dependent indicators of distraction

We used recent, behaviorally validated models to estimate the salience of individual distractors (Huang & Elhilali, 2017, 2020). This computed salience is, by definition, a proxy of the degree to which sound captures attention in a stimulus-driven fashion (Kaya & Elhilali, 2014). The finding that more salient sounds lead to stronger neurophysiological, sensory-driven responses may seem trivial at first glance. However, it is important to note that the specific features that determine the salience of a sound are still not fully understood. The fact that the computed salience predicted our data—derived from an entirely different experimental paradigm—confirms the generalizability of the salience model. Additionally, we observed a corresponding, salience-sensitive drop in target tracking, which suggests a distraction-induced attentional shift away from the target. Furthermore, attention-related modulation of sensory input begins at the brainstem level (Forte et al., 2017) and efferent signatures of attention have been found at the level of the cochlea (Gehmacher et al., 2022). While it is straightforward to extract voluntary attentional modulation experimentally, stimulus-driven attentional modulation is inevitably entangled with the properties of the sensory input. Eventually, it needs to be shown that the neurophysiological response mediates the effect of salience on behavioral performance.

While we failed to design a behavioral task that was sensitive to salience within subjects (performance around chance in surprise recall task), between subjects we found that the pupil response predicted both the drop in target tracking as well as target comprehension, both indicating a relationship between the pupil response and distraction. The absence of a direct effect of both distractor tracking and Δtarget tracking on target comprehension may be due to the combination of their low signal-to-noise ratio and the rather subtle manipulation of distraction during a high-context stimulus, which may have made target comprehension more robust to distraction.

Recent studies yield similar findings. In an EEG study by Holtze et al. (2021) subjects were listening to concurrent continuous speech while their own name was inserted into the to-be-ignored stream. In contrast to our findings, they observed that neural tracking of both the to-be-attended and the to-be-ignored streams increased after the name occurrence. While one’s own name is undoubtedly a very strong attention-capturing stimulus (Cherry, 1953; Moray, 1959), it is a unique example. Here we enhanced generalizability of results to other stimuli by using non-repetitive, real-world sounds of high spectro-temporal diversity. A similar approach was taken by Straetmans et al. (2021). Similar to our findings, they observed a drop in neural target tracking, both for the to-be-attended and to-be-ignored speech after the occurrence of a distractor sound. They also observed that the P3-component in the ERP to the distractor sounds (300 ms after distractor onset and thought to reflect attention orienting among others) was positively correlated with computed salience. Here we extend these results with pupillometry and instead of studying the event-related potential relative to sound onset, we examine the temporal response function in response to the sound envelope, which not only captures the initial onset of a sound but also its temporal modulation.

A large portion of the variance in the neurophysiological responses was left unexplained. On the one hand, this may be due to EEG and pupil size being inherently noisy signals, on the other hand, salience modelling may still have room for improvement (Duangudom & Anderson, 2007; Kalinli & Narayanan, 2007; Kaya & Elhilali, 2012; Kayser et al., 2005; Kim et al., 2014). Temporal and spatial context of a sound could be incorporated into salience modelling: for example, the sound of running water may not be very salient if you are out hiking, but would be very salient if you heard it in the walls of your home.

A further reason for substantial unexplained variance may be low variability in the predictors. By presenting all distractors at the same sound level, we minimized their variability in loudness, which is in the strongest determinant of salience in the model we used (Huang & Elhilali, 2017). The equalization may have resulted in a rather subtle manipulation of salience purely driven by their spectro-temporal content, such that the effect of acoustic salience on neurophysiological responses was underestimated. The fact that we still found an effect shows that current models of salience tap into aspects of sound beyond its level, that relate to stimulus-driven attention.

Subjective factors may also play a strong role here: for example, a cat may be more salient to someone who is afraid of cats than someone who has a neutral attitude towards cats. These individual aspects of salience may be captured in the statistical dependencies in the neurophysiological measures and their ability to predict behavior within-subjects. For example, a sound that is subjectively highly salient should both lead to strong neural tracking and pupil response, such that additional variance (beyond computed salience) in pupil response may be explained by distractor tracking. While we did not find additional connection between the neurophysiological responses on the within-subject level, on the between-subject level a larger pupil response led to a stronger drop in target tracking and worse comprehension. That means that the general distractibility of subjects could be inferred from their pupil response.

In sum, we provide evidence that stimulus-driven attention modulates neurophysiological responses during sustained attention to a target. Stimulus-driven attention is synonymous with distraction here, since the salient sounds were task-irrelevant but their presence had a consequence on behavioral performance (Makov et al., 2022). The neurophysiological measures allow us to index momentary distraction while subjects are focusing on a continuous stimulus. This may help to overcome challenges in the behavioral assessment of distraction: sustained attention does not need to be interrupted by a distractor-related task, which would make subjects perceive distractors as task relevant.

### Neurophysiological measures of distraction are orthogonal to hearing thresholds

We assumed that subjects with higher hearing thresholds would show enhanced distraction due to both less sharp attentional filters and fewer cognitive resources for attentional control. This should be reflected in stronger neurophysiological responses to distractors and worse comprehension even after compensation for hearing thresholds. We did not find a direct link between PTA and neurophysiological responses other than the drop in target tracking (which may be confounded, see above). But we found that higher PTA still led to worse comprehension. Within our design, it is not clear whether worse performance was due to distraction or if we would have received the same results with target speech in stationary noise only (i.e., without distractors), as might be expected, given general hearing loss. Still, we found neurophysiological responses to distractors that predict speech comprehension, such that we conclude that we captured neurophysiological markers which are orthogonal to hearing thresholds and explain variance beyond conventional measures of hearing loss such as PTA.

Finding measures that improve the explanation of inter-individual differences in speech-in-noise performance under high ecological validity are desirable for diagnostics and treatment of hearing challenges. Conventional speech-in-noise tests are performed in stationary noise, such that the factor of distractibility has not been considered to a greater extent, but it may be an important aspect in the design and fitting of hearing prosthesis. Furthermore, recent developments towards hybrid workplaces lead to the development of noise-adaptive hearing solutions (EPOS, 2023), which could benefit from objective measures of distraction.

## Conclusion

By relating neurophysiological responses evoked by task-irrelevant stimuli to both a current model for salience a well as behavioral performance, we provide evidence that neural tracking of distractor sounds, pupil response, and target tracking indicate momentary distraction. Hence, these measures may be suited to monitor the neural processing of distractors, which cannot be assessed behaviorally in such a direct fashion.

## Acknowledgements

EPOS Group A/S supported conference presentations. Torben Christiansen (EPOS Group A/S) contributed to the study design with discussion of potential application in noise-cancelling headsets. A first draft was proof-read by Bethany Plain.

## Supplemental material

**Figure 3-1:**
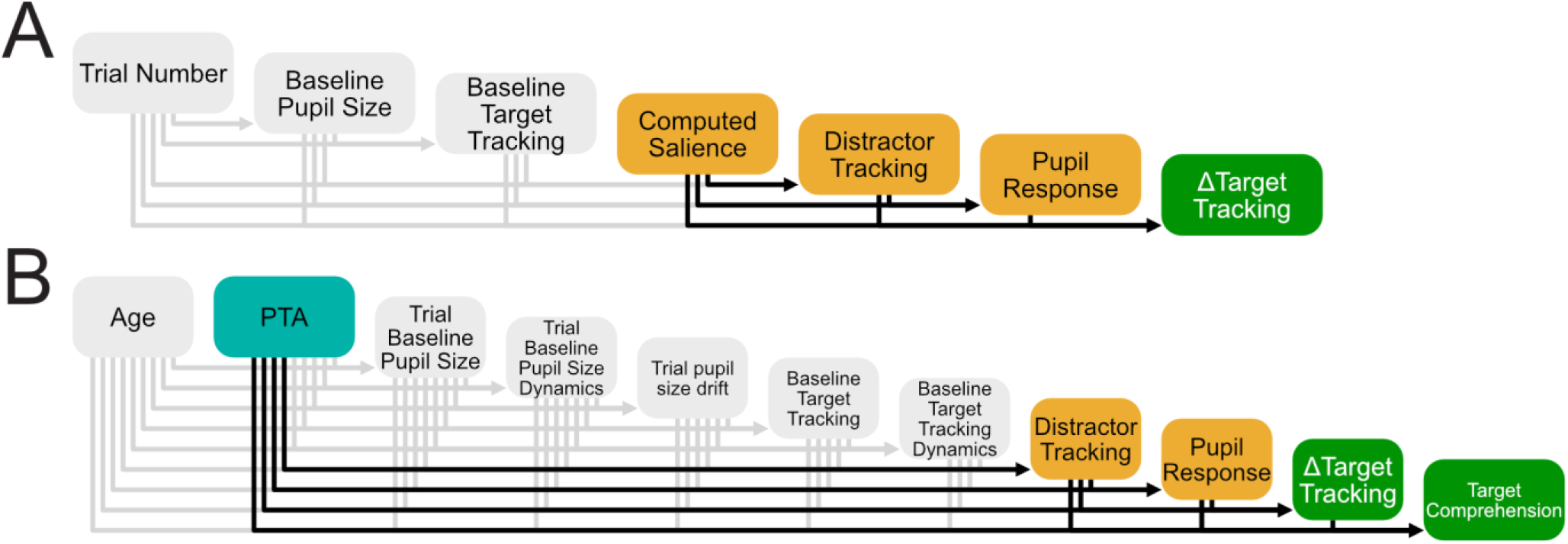
Hierarchical Statistical Modelling: **A)** Within-subject statistical modelling estimated the effect of stimulus features of single distractor sounds (computed salience) on single neurophysiological responses (distractor tracking, pupil response, Δtarget tracking) as well as their interdependence, while controlling for baseline measures (grey). **B)** Between-subject statistical modelling estimated the effect of hearing loss (PTA) on subjects’ average neurophysiological responses, and their common effect on target comprehension, while controlling for covariates (grey).

**Figure 4-1:**
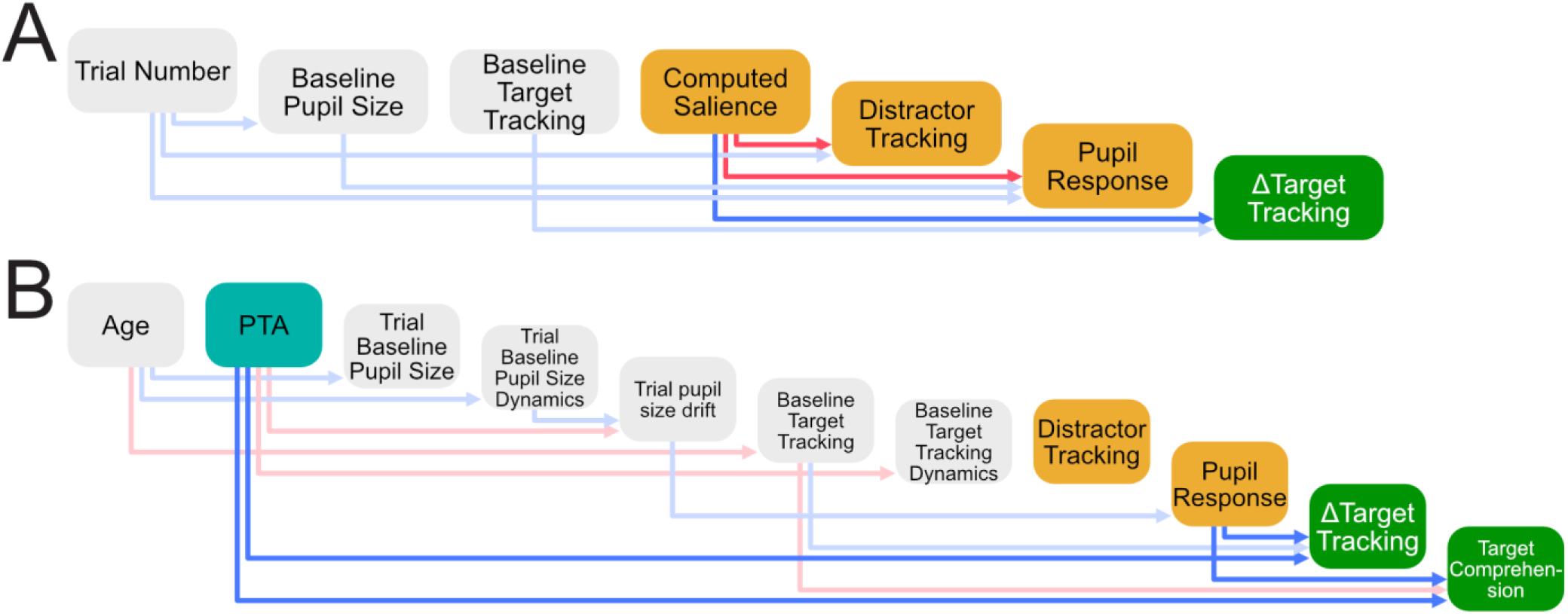
Results: Predictors among covariates (light arrows) and measures of interest (opaque arrows). Blue arrows indicate negative relationships, red arrows indicate positive relationships. Covariates are depicted in grey. **A)** Within-subject model. **B)** Between-subject model.

**Figure 6-1:**
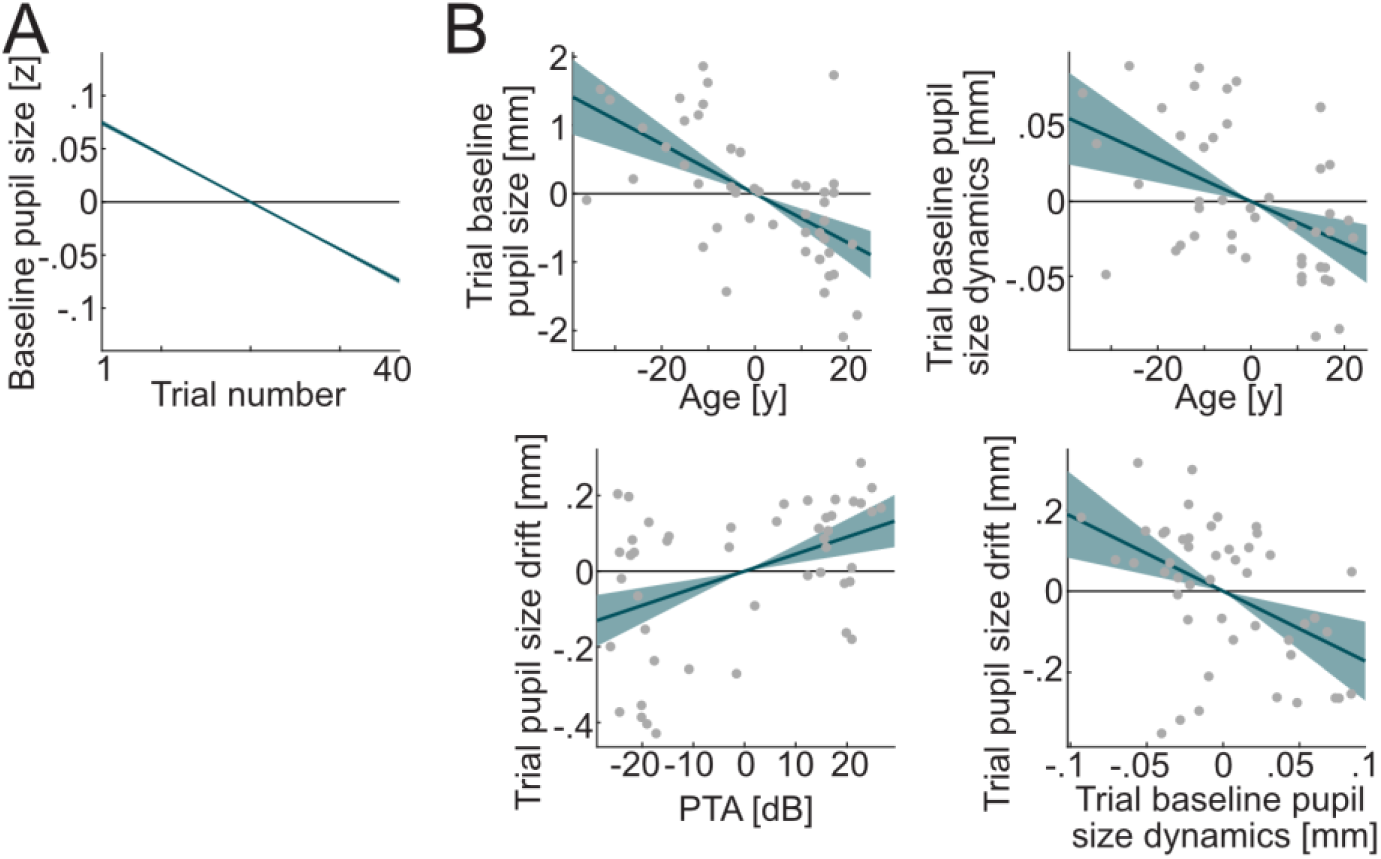
Covariates of pupil size. **A)** Within-subject estimates among pupil size covariates (all z-scored but trial number): Trial number (centered) on baseline pupil size. **B)** Between-subject estimates of predictors among pupil size covariates (all centered): Age on trial baseline pupil size, Age on trial baseline pupil size dynamics, PTA on trial pupil size drift, trial baseline pupil size dynamics on trial pupil size drift.

**Figure 7-1:**
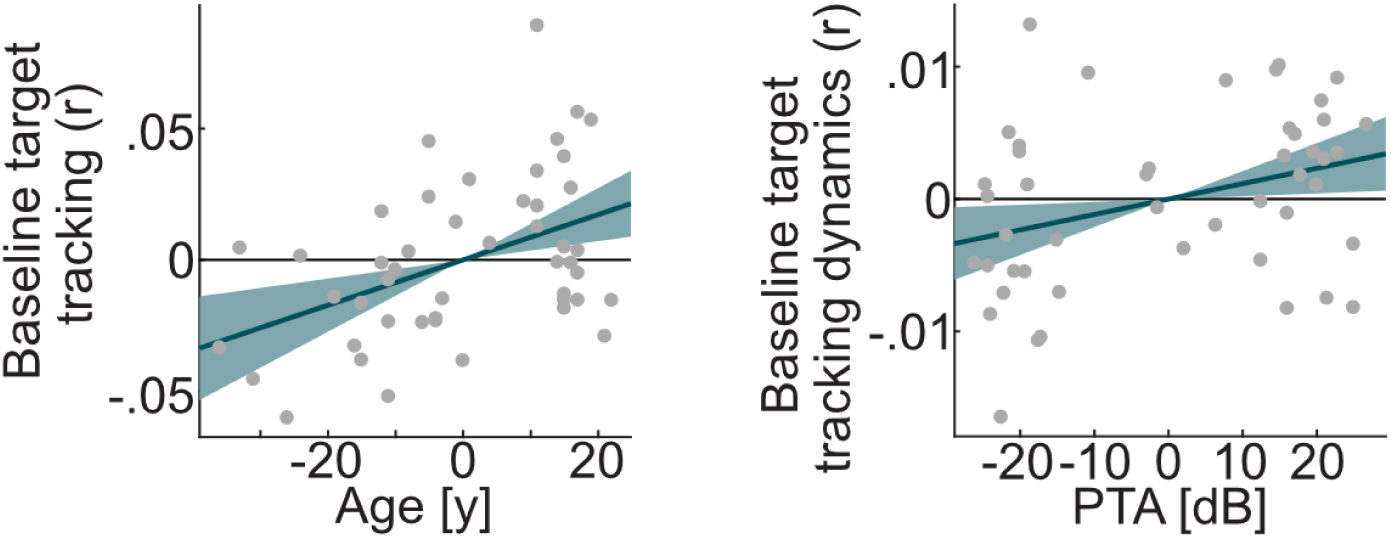
Covariates of target tracking. Between-subject estimates of predictors among pupil size covariates (all centered): Age on baseline target tracking, PTA on baseline target tracking.

